# Multiple inositol phosphate species enhance stability of active mTOR

**DOI:** 10.1101/2024.05.01.592113

**Authors:** Lucia E. Rameh, John D. York, Raymond D. Blind

## Abstract

Mechanistic Target of Rapamycin (mTOR) binds the small metabolite inositol hexakisphosphate (IP_6_) as shown in structures of mTOR, however it remains unclear if IP_6_, or any other inositol phosphate species, can activate mTOR kinase activity. Here, we show that multiple, exogenously added inositol phosphate species (IP_6_, IP_5_, IP_4_ and IP_3_) can all enhance the ability of mTOR and mTORC1 to auto-phosphorylate and incorporate radiolabeled phosphate into peptide substrates in *in vitro* kinase reactions. Although IP_6_ did not affect the apparent K_M_ of mTORC1 for ATP, monitoring kinase activity over longer reaction times showed increased product formation, suggesting inositol phosphates stabilize an active form of mTORC1 *in vitro*. The effects of IP_6_ on mTOR were reversible, suggesting IP_6_ bound to mTOR can be exchanged dynamically with the free solvent. Interestingly, we also observed that IP_6_ could alter mTOR solubility and electrophoretic mobility in SDS-PAGE in the presence of manganese, suggesting divalent cations may play a role in inositol phosphate regulation of mTOR. Together, these data suggest for the first time that multiple inositol phosphate species (IP_4_, IP_5_ and IP_6_) can dynamically regulate mTOR and mTORC1 by promoting a stable, active state of the kinase. Our data suggest that studies of the dynamics of inositol phosphate regulation of mTOR are well justified.

## Introduction

Inositol hexakisphosphate (IP_6_), also known as phytic acid or phytate, is a ubiquitous small metabolite found in many organisms, from yeast to mammals (Monserrate & York, 2010). IP_6_ and other inositol phosphate molecules comprise a family of signaling molecules derived from the cyclic polyhydroxy alcohol myo-inositol, which can be phosphorylated at six different positions of the inositol ring, generating six different species and several isomers of each for a total of 64 possible members, with 30 different species detected in cells (Irvine, 2005). IP_6_, the fully phosphorylated form of inositol phosphate is also the most abundant, with cellular concentrations in the range of 24-47 μM and blood levels around 30 μM, as recently confirmed by Capillary Electrophoresis coupled to Electrospray Ionization Mass Spectrometry (CE-EIMS) analysis (Qiu et al., 2020), and reaching 1 mM in plant seeds, which is equivalent to 1% of its dry weight (Irvine, 2005). IP_6_ can be further phosphorylated to generate the pyrophosphorylated forms IP_7_ and IP_8_ (Shears, 2016).

The role of IP_6_ in protein function is complex, with reports showing that this small molecule can serve as a competitive or allosteric regulator of enzyme activity, such as described for Casein Kinase 2 (CK2) (Lee et al., 2013), Bruton’ Tyrosine Kinase (Btk) (Wang et al., 2015), Yersinia outer-protein J (YopJ) (Zhang et al., 2016) and Histone Deacetylases (HDAC) (Watson et al., 2016), as a mediator of protein-protein binding to facilitate intra or inter-molecule complexes such as for Mixed Lineage Kinase Domain Like (MLKL) (Dovey et al., 2018) and Cullin/COP9 (Scherer et al., 2016) and protein oligomerization as for fibrinogen (Brehm et al., 2019). Furthermore, IP_6_ can function as a structural co-factor to promote proper protein folding. In fact, many X-ray crystallographic structures have shown unexpected electron densities consistent with IP_6_, present in the core of the protein (Blind, 2020). Some of them have been confirmed to be IP_6_ by mass spectrometry. As a structural co-factor, IP_6_ association with its protein target must be tight and long-lived and must be buried within the core of the protein, as described for ADAR2 (Macbeth et al., 2005)

Although less abundant than IP_6_, other inositol phosphate species have been shown to play critical roles in cell signaling as well. For example, IP_3_ (inositol-1,4,5-P_3_) is well known for its role as a second messenger for various growth factor signals that regulate intracellular calcium release and IP_4_ (inositol-1,4,5,6-P_4_) was recently shown to allosterically regulate histone deacetylases (HDAC), through direct binding (Maffucci & Falasca, 2020). Some of the roles of inositol phosphates in protein regulation are specific to a certain species and/or isomer while others are shared between multiple species.

Cryo-EM studies and retrospective analysis of crystallographic structures revealed that IP_6_ co-crystalizes with mTOR (Gat et al., 2019; Scaiola et al., 2020), a serine/threonine kinase which is the core catalytic subunit of two complexes, mTORC1 and mTORC2. While both complexes require the accessory protein mLST8 (mammalian homolog of protein Lethal with Sec Thirteen), mTORC1 is characterized by the regulatory subunit Raptor and mTORC2 by Rictor. mTORC1 is a nutrient-sensing kinase that signals for increase in anabolic processes in times of nutrient abundance (Mossmann et al., 2018; Saxton & Sabatini, 2017). The combination of growth factors and nutrients, such as amino acids, lipids and glucose, activates mTORC1 at the lysosomal surface by promoting translocation of mTORC1 from the cytosol to the lysosomal surface and activation of lysosomal RHEB (RHEB/GTP), an allosteric activator of mTORC1 (Yang et al., 2017). However, RHEB-independent mechanisms for mTORC1 activation must exist, especially considering that mTOR and its substrates are found in different organelles and subcellular locations. In fact, phosphatidic acid was recently shown to activate mTOR in the absence of RHEB (Frias et al., 2023). To this date, there are no soluble small metabolite shown to directly regulate mTOR catalytic activity.

Interestingly, IP_6_ was found within a highly positive pocket formed by the FAT domain of mTOR referred to as the I-site (Scaiola et al., 2020). The FAT domain of mTOR forms a C-shaped solenoid structure that surrounds the N- and C-lobes of the kinase domain and was shown to participate in RHEB/GTP induced conformational changes that culminate in kinase activation (Chao & Avruch, 2019). Mutation of two or three of the residues that coordinate IP_6_ binding inside the FAT domain were used to address the role of IP_6_ in mTOR complex formation and activity. In one study, mutation of two lysines within the I-site (K1753/1788E) abolished *in vitro* kinase activity of truncated mTOR (Gat et al., 2019), whereas in another study in which mTOR activity was measured in the presence of its regulatory subunits (LST8, Rictor and SIN1) the same mutations had no impact on complex formation or kinase activity (Scaiola et al., 2020). These seemingly contradictory results led Scaiola and co-investigators to conclude that IP_6_ binding to mTOR is dispensable for kinase function when the regulatory subunits are present. Both groups proposed that IP_6_ plays a structural role by allowing proper folding of the kinase domain. Surprisingly, neither study addressed whether exogenous IP_6_ can enhance mTOR kinase activity and whether this role of IP_6_ can be fulfilled by other inositol phosphate species.

Here, we examined the impact of various inositol phosphate species on mTOR kinase activity and stability/solubility, in the context of mTOR alone or complexed with its regulatory subunits LST8 and Raptor (mTORC1). The data suggest that exogenous IP_6_, IP_5_, IP_4_, and IP_3_ (to a lesser extent) enhanced mTOR and mTOR/LST8/Raptor autophosphorylation or phosphorylation of exogenous peptide substrates in a dose-dependent and saturable manner. This could partially be explained by an increase in solubility/stability of the enzyme. The effect of IP_6_ on mTOR was dynamic and prolonged the active state of the enzyme without affecting the apparent K_M_ for ATP or the peptide substrate.

## RESULTS

### Higher order inositol phosphates increase auto- and peptide-phosphorylation by mTOR and mTOR/LST8/Raptor

To better understand how IP_6_ and other inositol phosphate species alter mTOR catalytic activity, we measured mTOR autokinase using radiolabeled-ATP in the presence or absence of exogenous 100 μM inositol or inositol phosphate species during 1hr incubation time. Radiolabeled phosphate incorporation into both mTOR and mTOR/LST8/Raptor were enhanced several-fold by exogenous IP_4_, IP_5_ or IP_6_, but not by inositol, IP_1_ or IP_2_ **(Fig 1A-B),** despite equal amount of protein being added to all these reactions. IP_3_ also enhanced mTOR and mTOR/LST8/Raptor autophosphorylation although to a lesser extent than the higher phosphorylated forms of inositol.

**Figure 1:**
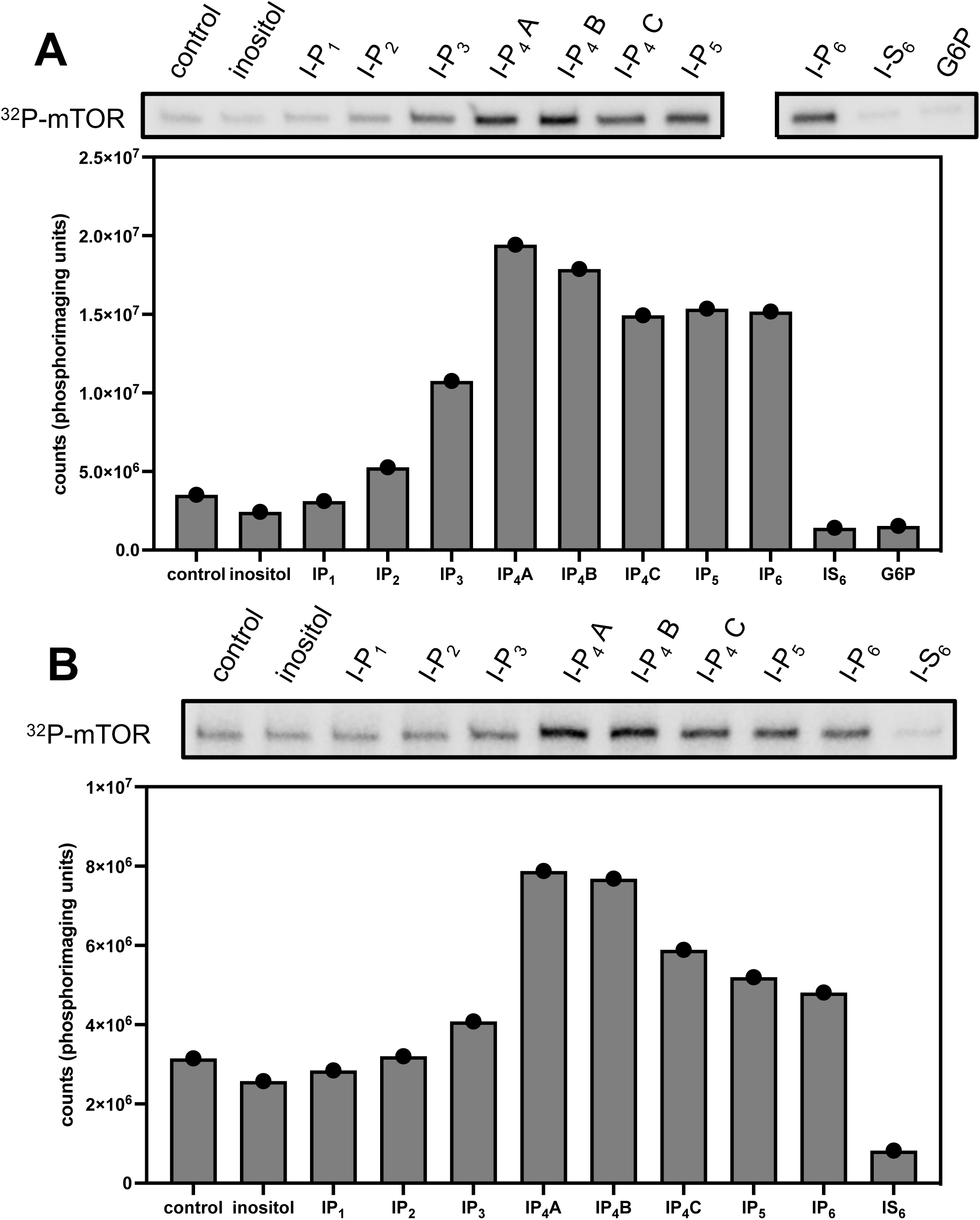
Inositol phosphates increase autophosphorylation of mTOR when assay alone (A) or in the in the presence of LST8 and Raptor (B). mTOR or mTOR/LST8/Raptor were incubated with [^32^P] γ-ATP in kinase reaction without (control) or with 100 μM of inositol or various inositol phosphate species, IS_6_ or glucose-6-phosphate (G6P), as indicated. Reactions were stopped with EDTA before addition of SDS-loading buffer for electrophoresis (see section 4). Shown are the phosphoimager images (upper panels) and quantification (lower panels). Coomassie staining of the gels are shown in Supplemental Figure S1.

Three different isoforms of IP_4_ were tested with similar outcomes **(Fig. 1)**. As a control, we used glucose-6-phosphate (G6P) and inositol hexa-kis-sulphate (IS_6_) which like IP_6_ is highly negatively charged. However, IS_6_ was unable to enhance phosphate-labeled mTOR and rather inhibited it **(Fig. 1)**, showing that the effect of IP_6_ on mTOR was not mimicked by another highly negatively charged small molecule. Finally, we noted that inositol phosphate-dependent increase in auto-phosphorylation was more robust in mTOR alone (about 6-fold) than in mTOR co-expressed with LST8 and Raptor (about 2 to 3-fold). These data suggest that multiple inositol phosphate species enhance mTOR and mTORC1 autokinase in *in vitro* reactions.

To more thoroughly characterize the effect of inositol phosphates on mTOR, we measured the phosphorylation of peptide substrates in the presence of various concentrations of inositol phosphates, again using radiolabeled ATP incorporation during 1-2 hrs assays. We used a peptide substrate derived from the mTORC1 phosphorylation site in 4EBP, which has been successfully used in other studies of mTOR catalytic activity (Kang et al., 2013). IP_5_ and IP_6_ at 1 μM maximally increased peptide phosphorylation by mTOR or by mTOR/LST8/Raptor as compared to vehicle control **(Fig. 2A-B)**. At 0.1 μM, IP_6_ effect was submaximal, showing a concentration-dependent effect **(Fig. 2C)**. We calculated through non-linear curve fit an EC50 between 0.07 and 0.17 μM for the IP_6_ effect on mTOR. IP_4_ enhanced substrate phosphorylation to similar extent as IP_5_ and IP_6_, but higher concentrations were required (EC50 between 0.44 and 2.4 μM), and at least 10 μM for maximal activity **(Fig. 2 A-C)**. The effect IP_3_ on substrate phosphorylation by mTOR and mTOR/LST8/Raptor was submaximal even at 100 μM **(Fig 2 A-B)**. These data suggest that multiple inositol phosphate species enhance mTOR and mTORC1 peptide phosphorylation in equilibrium reactions.

**Figure 2:**
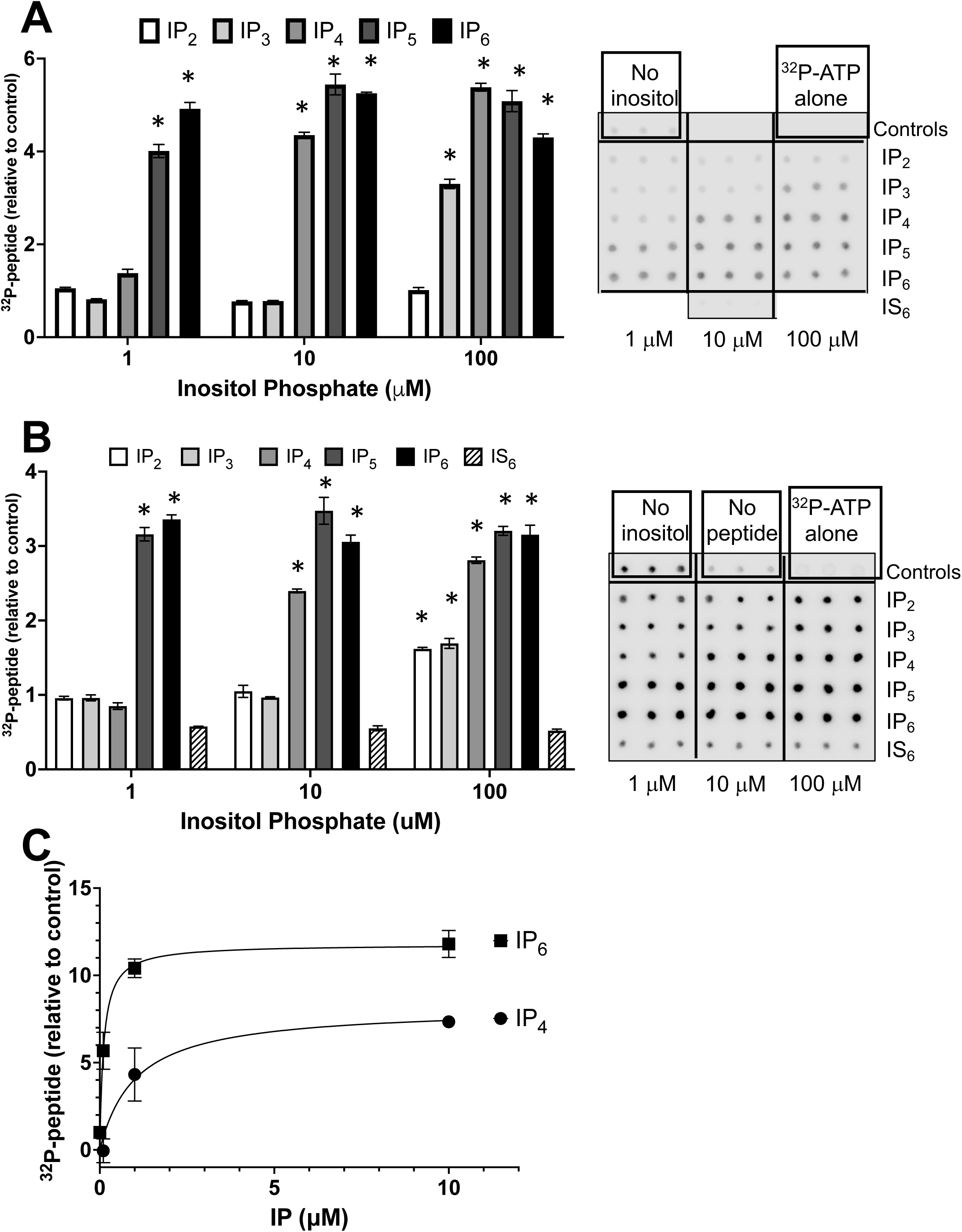
Inositol phosphate species enhance mTOR phosphorylation of peptide substrate with different affinities. mTOR kinase assay using 4EBP peptide as a substrate and recombinant mTOR alone (A and C) or mTOR/LST8/Raptor (B). mTOR or mTOR/LST8/Raptor were incubated with [^32^P] γ-ATP in kinase reaction without (control) or with 1, 10 or 100 μM of various inositol phosphate species (see materials and methods for isomers used), inositol IS_6_, as indicated. Reactions were stopped with EDTA before spotting on P81 paper. (C) shows Michaelis-Menten-style plot for product generated as a function of concentration of exogenous inositol phosphate (from data shown in Fig S2). (A and B) Right panels show phosphorimager images of the spotted P81 papers and left panels show mean and standard deviations of quantified triplicate spots. (*) indicate statistically significant changes (P<0.0001) using student’s t-test as compared to control (control=1).

Since IP_6_ had been previously shown to inhibit serine-threonine phosphatases type 1, 2A and 3 (Larsson et al., 1997) with K_M_s around 10μM, we tested whether the increase in peptide phosphorylation in our assays could be due to decreased phosphatase activity (which could potentially co-purify with mTOR) by IP_6_. If this was true, we expected to see blunting of the IP_6_ effect when broad spectrum phosphatase inhibitors were present. Addition of NaFl, ß-glycerophosphate and nor-cantharidin to the mTOR kinase reaction had little effect on the amount of phospho-peptide detected and did not mask the IP_6_ effect **(Fig. 3)**, suggesting that the observed IP_6_ effect is most likely independent of any co-purifying phosphatase activity in the reactions.

**Figure 3:**
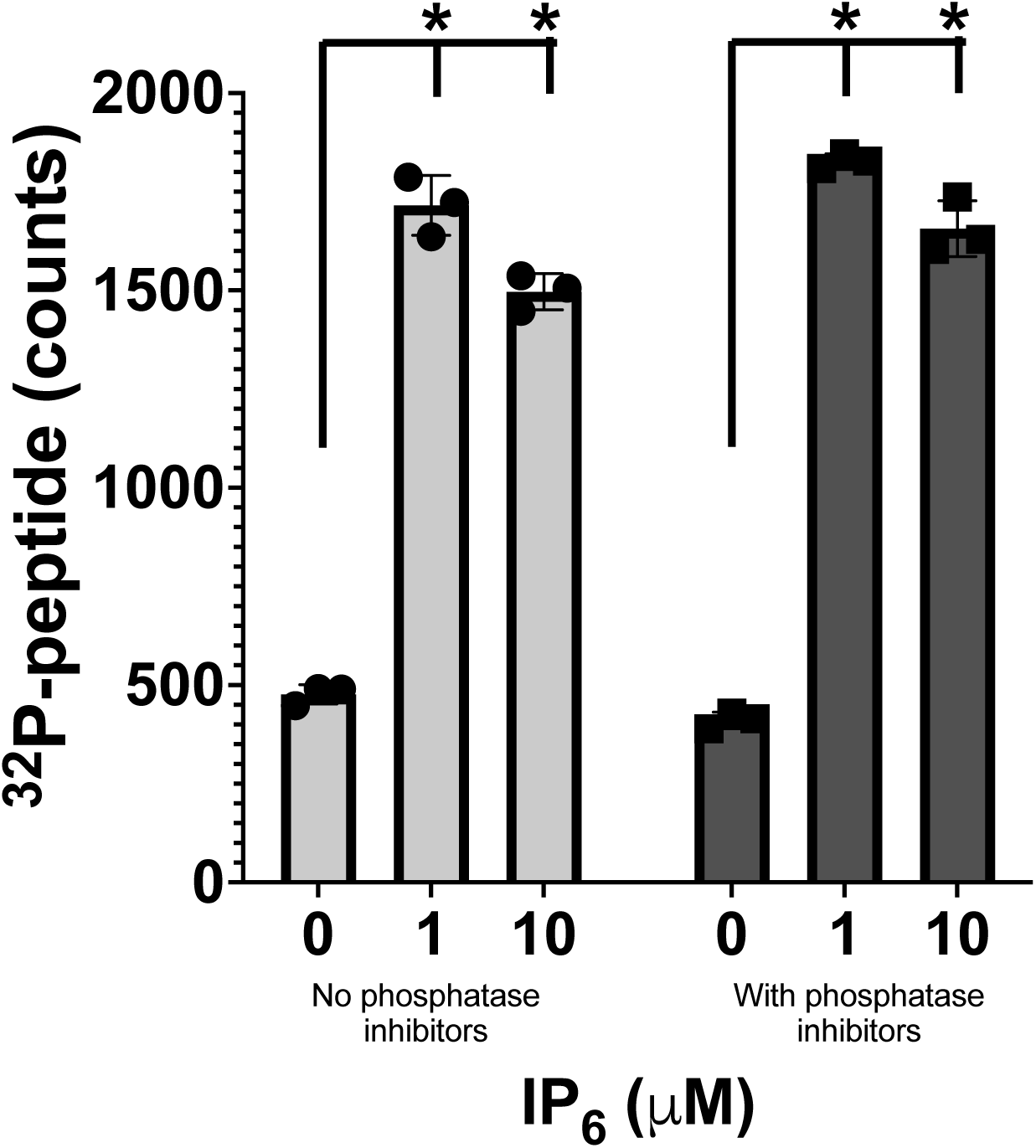
IP_6_ effect on peptide phosphorylation by mTOR is not affected by phosphatase inhibition. mTOR was incubated with [^32^P] γ-ATP in kinase reaction without (control) or with 1 or 10 μM of IP_6_ in kinase buffer supplemented or not with phosphatase inhibitors, as indicated. Data shown is the mean and standard deviations of quantified triplicate spots. (*) indicate statistically significant changes (P< 0.00001) using student’s t-test as compared to control.

Thus, sub-physiological concentrations of inositol phosphates enhanced mTOR’s or mTOR/LST8/Raptor’s ability to phosphorylate exogenous or intramolecular substrates during *in vitro* assays. The sensitivity of mTOR to exogenous inositol phosphate species was directly proportional to the number of phosphates in the molecule (IP_6_= IP_5_ >IP_4_>IP_3_>IP_2_) but neither G6P nor IS_6_ had any detectable effect.

### IP_6_ increases mTOR solubility in the presence of manganese

We noticed that in the absence of inositol phosphates, mTOR activity plateaued at around 10 to 15 minutes **(Supplemental Fig S3)**. We did not suspect that mTOR was being proteolytically cleaved because no fragments of mTOR were observed after one-hour incubations **(Supplemental Fig S1)** and our assays regularly contained protease inhibitors. Thus, we suspected that solubility of mTOR could decrease over time. To examine this possibility, we collected one quarter of the supernatant of the kinase reactions at different incubation times. mTOR solubility decreased rapidly over time with 68% (17/25) of mTOR being soluble at time zero (samples on ice) down to only 16% (4/25) soluble after 30 minutes at 30°C **(Fig 4A)**. When IP_6_ was added to the reaction, 64% (16/25) of mTOR was still in solution after 30 minutes, which is a four-fold increase as compared to control.

**Figure 4:**
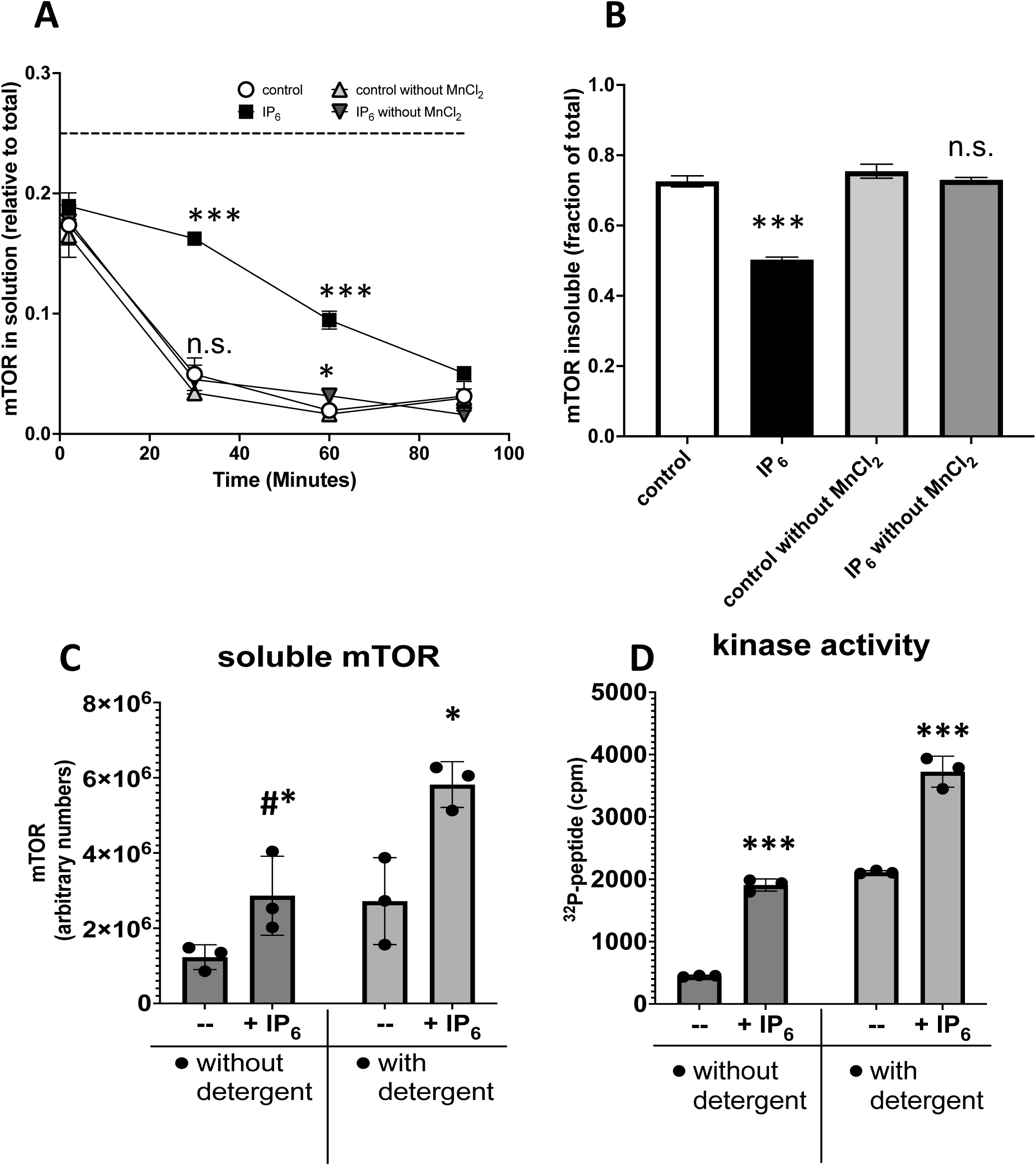
IP_6_ increases solubility of mTOR. mTOR kinase reactions with or without IP_6_ (1 μM) and with or without MnCl_2_ (10 mM) were sampled over time and analyzed by western blot for mTOR (soluble fraction), followed by a final extraction of the material left in the tube after 90 minutes (insoluble). In A, the fraction of soluble mTOR (soluble/ total) over time of incubation was plotted. Dash line indicates the maximal possible amount of mTOR if completely in solution (0.25) given the volume collected (1/4). In B, the fraction of insoluble mTOR (insoluble/total) at the end of the reaction was plotted. In C and D, mTOR kinase was assayed with and without 0.1% CHAPS and with or without IP_6_ (10 μM) and samples collected at the end of 90 minutes for western-blot of mTOR (C) or [^32^P]-peptide analysis (D). Results shown are the mean and standard deviation of triplicate samples. (*) indicates that IP_6_ changes are statistically significant as compared to equivalent control without IP_6_ with (A and B) (*) P=0.017; (***) P<0.0002; n.s= non-significant; (C and D) (***)P<0.0037; (#*)P=0.062; (*) P=0.015.

Concomitant with the increase in IP_6_-dependent mTOR solubility, we observed a decrease in the mTOR fractions that were out of solution and only extractable by SDS/boiling **(Fig 4B)**. Interestingly, when MnCl_2_ was omitted from the reactions, the IP_6_ effect was negligible, with only 18% (4.5/25) in solution after 30 minutes at 30°C. IP_6_ forms a high affinity complex with divalent cations, including manganese (Torres et al., 2005). It is possible that MnCl_2_ increases the affinity of IP_6_ for mTOR due to the ability of manganese to shield negative charges of IP_6_, similar to the effect of Mn^2+^/Mg^2+^ on ATP binding to kinases, however this remains to be tested. We next compared the effect of IP_6_ with CHAPS, a detergent that is commonly used to solubilize mTOR *in vitro*. IP_6_ at 10 μM had a similar effect as 0.1% CHAPS by doubling the fraction of mTOR in solution **(Fig 4C)**. Both, CHAPS and IP_6_ increased mTOR kinase to the same extent **(Fig 4D)**. Interestingly, IP_6_ further enhanced mTOR solubility and activity in the presence of CHAPS, which is present at 100-fold higher molar concentration than IP_6_, consistent with IP_6_ and CHAPS contributing to mTOR solubility in different ways. It is not clear whether the insoluble mTOR is completely inactive or whether it can contribute to kinase activity to some extent. These data suggest that IP_6_ increases mTOR solubility.

### IP_6_ prolongs mTOR/LST8/Raptor active state without affecting affinity for ATP

mTORC1 had higher stability than mTOR alone, with 90-100% retention of mTORC1 activity (peptide phosphorylation during 1 hr incubation) in the sample after 2 hours incubation at room temperature (**Supplemental Fig S4**). These observations are consistent with the role for mTORC1 regulatory subunits in stabilizing the complex. Due to the higher stability of mTORC1, we were able to measure initial velocity of the enzyme with or without exogenous IP_6_ and with varying concentrations of ATP (**Supplemental Fig S5**). Michaelis-Menten plots revealed that IP_6_ had no effect on mTORC1 apparent K_M_ for ATP (**Fig 5A-B**), which was within the expected range of 50μM, as previously reported (Kang et al., 2013). In fact, we calculated that the relative K_M_ for ATP was slightly higher in mTORC1/ IP_6_ than mTORC1 alone. When kinase reactions were carried out at equilibrium (4-16 hrs), there was an increase in product formation by mTORC1 when IP_6_ was present, consistent with IP_6_ stabilizing an active or soluble form of the kinase. We have calculated that less than 10% of the peptide substrate is phosphorylated even during extended reactions, making it unlikely that substrate availability is limiting. We also measured velocities with varying concentrations of peptide substrate during 0.5 to 2-hour reactions (**Fig 6A-B**). Michaelis-Menten plots of velocity over substrate concentration showed almost identical curves whether IP_6_ was present or not **(Fig 6C)**. Measurements of substrate dependence during equilibrium reactions (8-10 hrs) showed an increase in phospho-peptide when IP_6_ was present, an effect that was seen regardless of the peptide substrate concentration **(Fig 6D)**. Together, these results suggest that IP_6_ preserves mTORC1 in an active and/or more soluble state for longer times during the kinase reactions, without affecting the K_M_ for ATP of peptide substrate.

**Figure 5:**
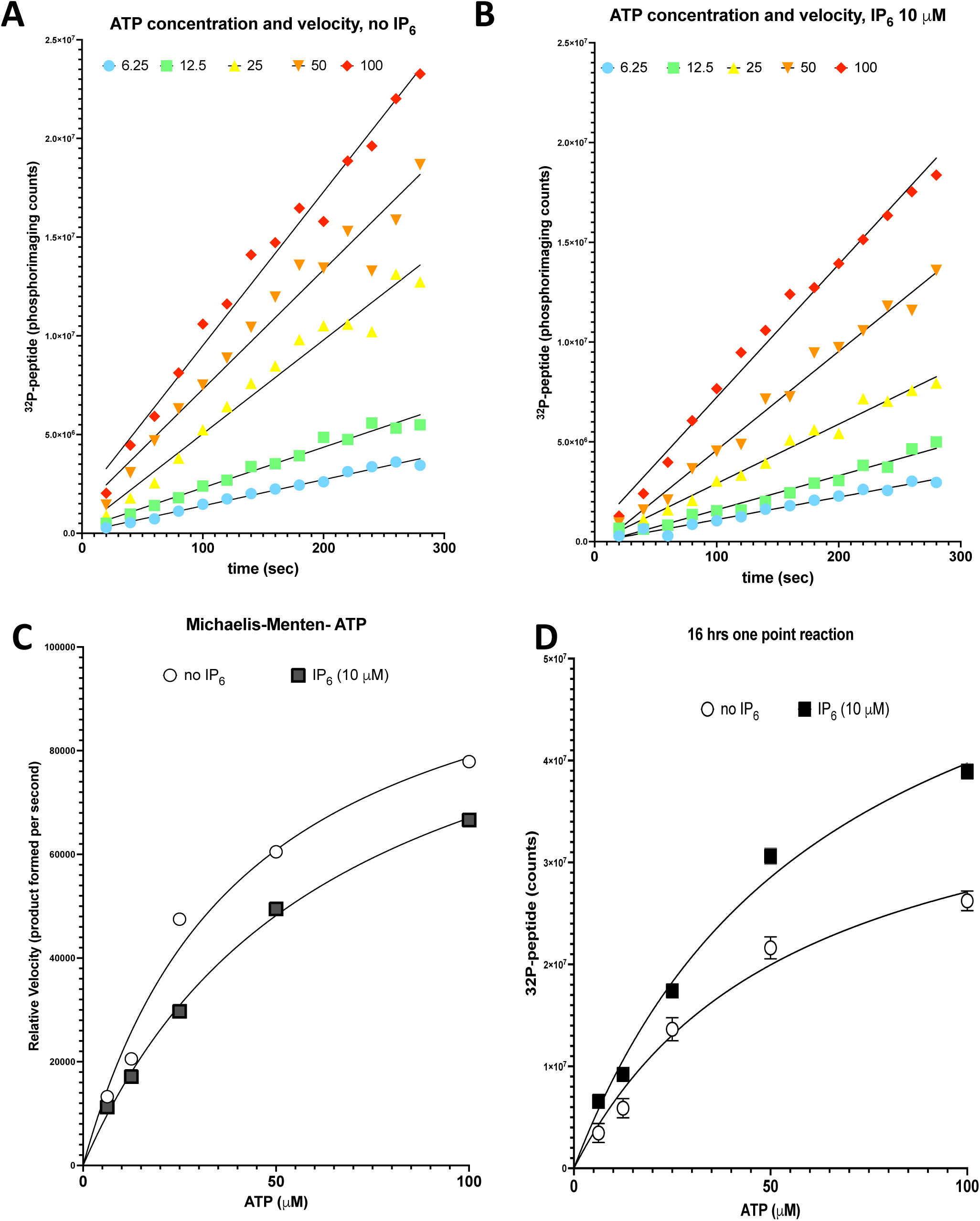
IP_6_ does not affect mTOR/LST8/Raptor affinity for ATP but it increases product accumulation during prolonged incubation. Initial velocities of mTORC1 in the absence (A) or presence (B) of IP_6_ (10 μM) and varying concentrations of [^32^P]-ATP/ATP. Michaelis-Menten plots of initial velocity (product in counts/second) as a function of ATP concentration (C). Non-linear regression curve fit shows a relative K_M_ for ATP of 41.5 and 64 μM for reactions without or with IP_6_, respectively and V_max_ of 1.11×10^5^ and 1.09 x10^5^ (counts/sec), respectively. Product formed over prolonged incubation (16 hrs) as a function of ATP concentration (D). From non-linear regression curve fit, we extrapolate that mTOR required ATP at 56 μM (without IP_6_) and 64 μM (with IP_6_) for half of the maximal product formation and that the maximal product formed with IP_6_ was 1.54-fold higher than without IP_6_.

**Figure 6:**
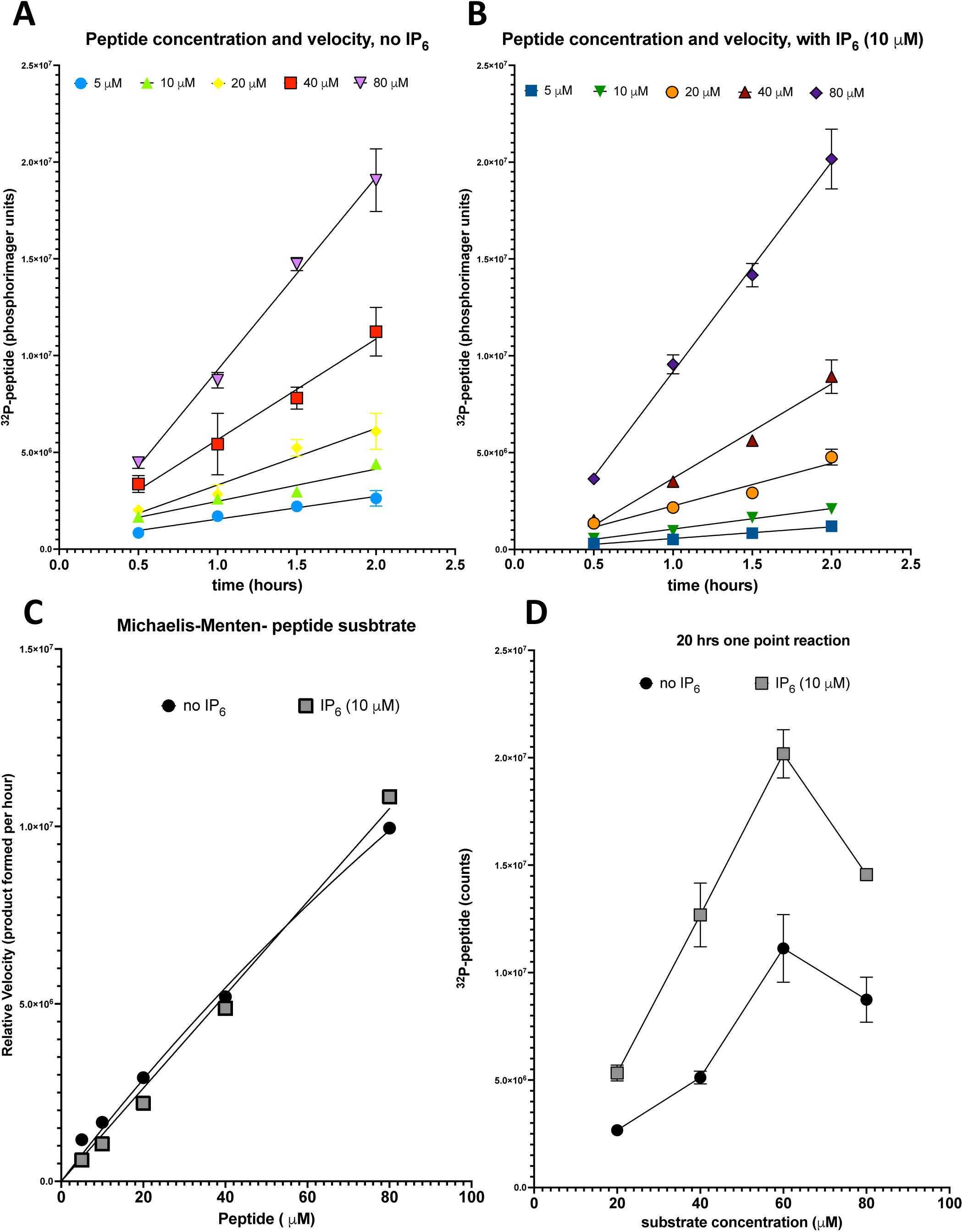
IP_6_ does not affect mTOR/LST8/Raptor affinity for peptide substrate but it increases product accumulation during prolonged incubation. Velocities of mTOR/LST8/Raptor over 2 hrs incubation in the absence (A) or presence (B) of IP_6_ (10 μM) and varying concentrations of 4EBP peptide substrate. Michaelis-Menten plots of initial velocity (product in counts/second) as a function of peptide substrate concentration (C). Calculations of relative K_M_ were not possible because V_max_ was not obtained. Product formed over prolonged incubation (18 hrs) as a function of peptide substrate concentration (D).

### Association of inositol phosphates with mTOR in the presence of manganese promotes an electrophoretic mobility shift

While examining the effect of IP_6_ on mTOR or mTOR/LST8/Raptor autokinase, we noticed that when exogenous IP_6_ was added to the reaction, mTOR electrophoretic mobility shifted in denaturing SDS-PAGE, to an apparent molecular weight band of about 100 K_Da_ higher than expected for the recombinant mTOR in SDS-PAGE, which is around 137 K_Da_ (**Fig 7A**). Using western blotting we confirmed that this IP_6_-dependent higher shifted band cross-reacted with mTOR antibodies (**Fig 7B**). The formation of this super-shifted mTOR band was IP_6_-concentration-dependent (**Fig 7C-E**) and was observed with IP_6_ concentrations as low as 1μM (**Fig 7E**), which is similar to the dose-response observed for kinase assays. ATP was not required for the IP_6_-dependent mTOR electrophoretic mobility shift, and addition of the mTOR kinase inhibitor KU0063794 did not prevent it (**Fig 7D**). When added after heat-induced denaturation of mTOR, the effect of IP_6_ on mTOR electrophoretic mobility was lost, indicating that proper mTOR folding was required for the IP_6_ super-shift effect (**Fig 7B**).

**Figure 7:**
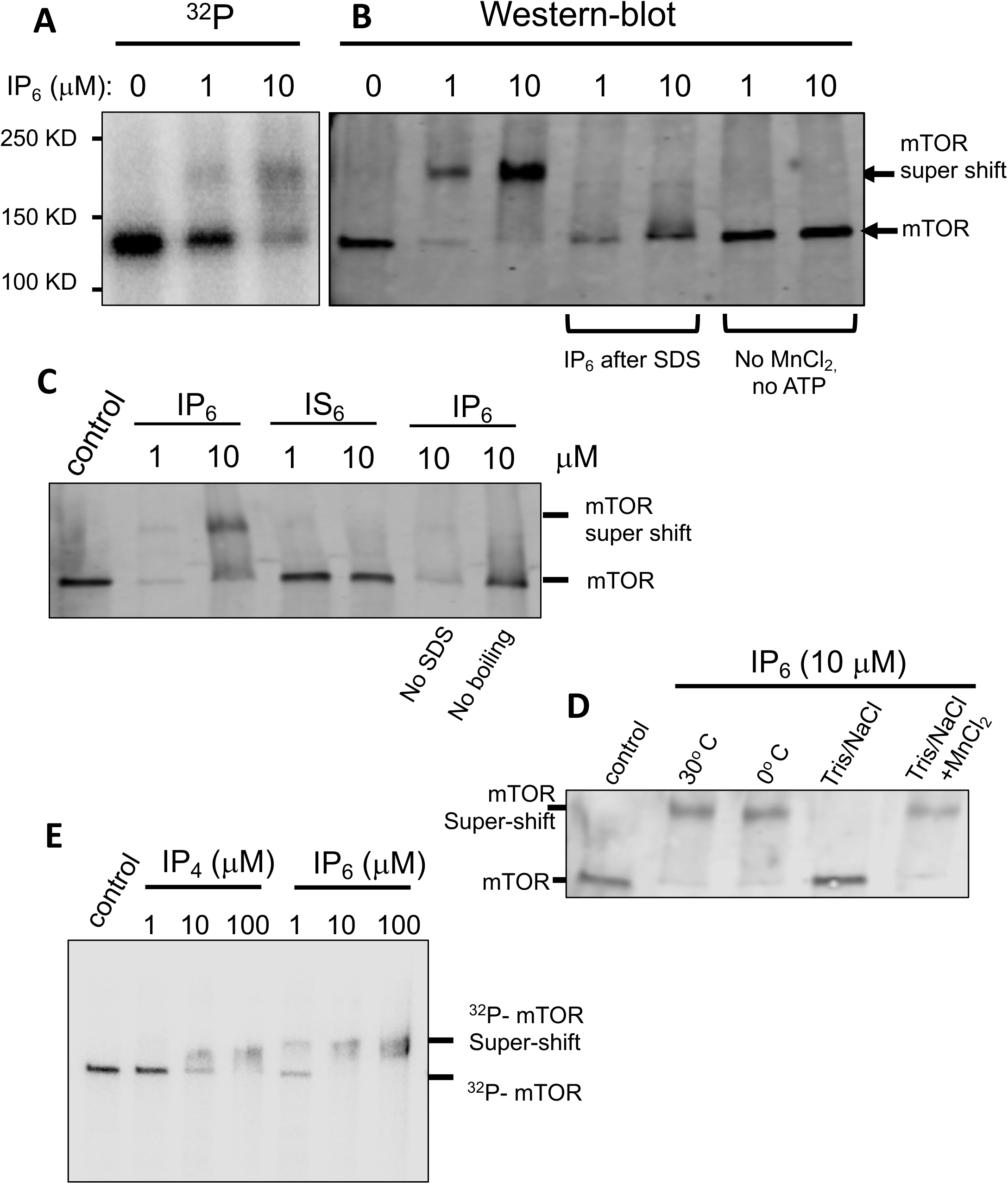
IP_4_ and IP_6_, but not IS6, promote the formation of mTOR super-shift in electrophoretic mobility in a dose-dependent manner. Panels A and E are phosphorimager images of mTOR after autokinase reaction with [^32^P]-ATP, without (control) or with IP_6_ or IP_4_ (E) at the concentration indicated. Panels B, C and D show western-blots of mTOR after autokinase reaction with unlabeled ATP and without (control) or with IP_6_ or IS_6_ (C), as indicated. In panel B, lanes 4 and 5, IP_6_ was absent at the kinase reaction and added after SDS-loading buffer. In 5 and 7, ATP and MnCl_2_ were not added to the kinase reaction. In panel C lanes 6, loading buffer without SDS was used and in lane 7, samples were not boiled at 100°C. In panel D, lane 3, kinase reaction was kept at 0°C; lane 4, reaction was carried out with Tris NaCl buffer; lane 5 is similar to lane 4 except that MnCl_2_ at 10 mM was present. All reactions were stopped with loading buffer and without the addition of EDTA.

Furthermore, in the absence of MnCl_2_, the effect of IP_6_ to induce the shift was abolished (**Fig 7B** and **Fig 7D**). It is notable that this MnCl_2_ requirement was also observed for IP_6_’s effect on mTOR solubility (**Fig 4A**), and that heating the proteins at 100°C prior to SDS-PAGE as part of the denaturation of the samples was necessary for the formation of this super-shift, suggesting that the IP_6_ modification of mTOR is denaturation-resistant (**Fig 7C**). Addition of EDTA after IP_6_ incubation with mTOR and before boiling abolished the formation of the super-shift (SDS-PAGE in **Fig 1** have EDTA added), confirming that metal ions are important not only during incubation but also during the heating step to observe this shift. IS_6_ was unable to drive the formation of this high molecular weight band (**Fig 7C**), whereas IP_4_ was able to induce the super shift in a concentration-dependent manner (**Fig 7E**), as for the kinase assays. Thus, the ability of inositol phosphates to promote mTOR super-shift positively associates with mTOR’s ability to phosphorylate its substrates. We propose that mTOR super-shift can be used as a non-radiometric assay of inositol phosphate binding to mTOR.

### IP_6_ activation of mTOR is dynamic

IP_6_ is found associated with several proteins and it is thought to function as a structural co-factor to drive proper protein folding. In mTOR, IP_6_ binds to the “I-site”, a pocket within the FAT domain that contains 5 positively-charged residues(Scaiola et al., 2020). When these residues were all mutated into negatively-charged residues, the protein could not be expressed, supporting the idea that the I-site is necessary for proper mTOR protein folding (Gat et al., 2019; Scaiola et al., 2020). Intriguingly, our data show that exogenous IP_6_ had a robust effect on the solubility and activity of mTOR expressed in insect cells. If IP_6_ is indeed a structural co-factor for mTOR, we expected that the recombinant mTOR protein used in our assays would co-purify with IP_6_, since insect cells have abundant IP_6_. Instead, the data suggested that IP_6_ association with mTOR may be more dynamic than previously appreciated. To test if IP_6_ binding to mTOR is long-lasting, we pre-incubated mTOR with buffer containing MnCl_2_ and either 0.1 or 1.0 μM IP_6_, which are concentrations that promote sub-maximal or maximal activation of mTOR, respectively (**Fig 2C and S2**). Each of these were diluted and assayed for phosphate incorporation into peptide substrate in reactions that had a final concentration of either 1.0 or 0.1 μM IP_6_. When mTOR was pre-incubated with IP_6_ at 0.1 μM and kept at 0.1 μM, the kinase reactions were initially slow and plateaued at about 15 minutes (**Fig 8A**), in contrast to the reactions in which mTOR was assayed with 1μM IP_6_, which were initially faster and plateau at 20 minutes (**Fig 8A**). On the other hand, when mTOR was pre-incubated with 1.0μM IP_6_ but then switched to lower 0.1μM, the kinase reactions were initially faster but then slowed after 10 minutes and plateaued together with the mTOR that was pre-incubated and kept at 0.1μM (**Fig 8A-C**). These data suggest IP_6_ has the capacity to dissociate from mTOR when diluted out, consistent with mTOR dynamically “sensing” the IP_6_ levels in the kinase reactions.

**Figure 8:**
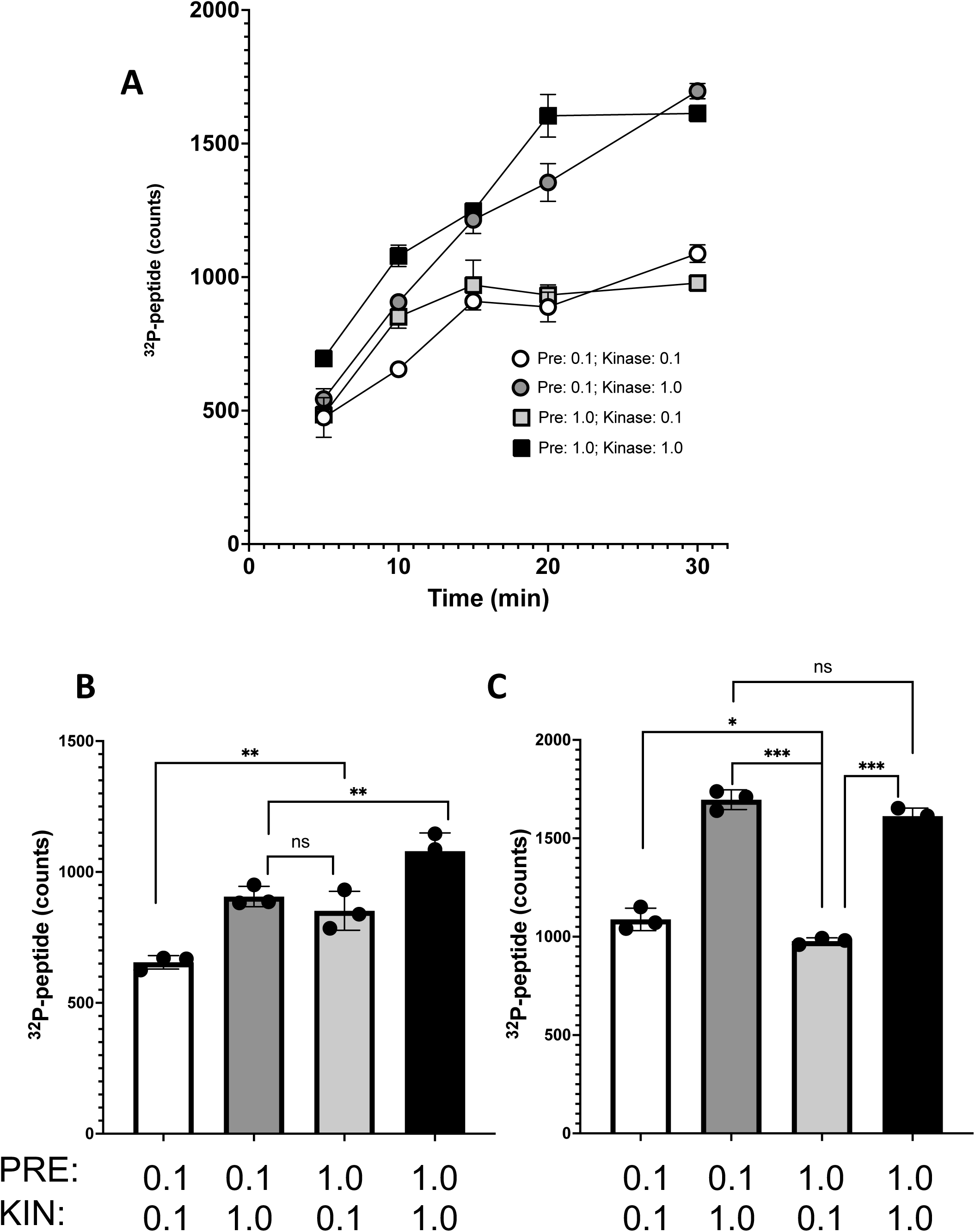
The IP_6_ effect on mTOR kinase is short-lived. Peptide phosphorylation over time in kinase assays in which mTOR was pre-incubated with either low IP_6_ (0.1 μM) or high IP_6_ (1.0 μM) together with MnCl_2_ (10 mM). After pre-incubation, mTOR was diluted into complete kinase buffer containing either low IP_6_ (0.1 μM) or high IP_6_ (1.0 μM), as indicated. Samples of the reactions were collected at the time point indicated and analyzed for [^32^P]-peptide. Results shown are the mean and standard error of 3 samples. In A, all time points are shown. In B and C, the bars represent the data from the 10 and 30-minute samples, respectively. (*) indicates statistically significant changes as determined by a paired student’s t-test with (*) P=0.032; (**) P<0.02; (***) P<0.0001.

Together, these data provide the first comprehensive analyses of inositol phosphate regulation of mTOR, suggesting 1. multiple inositol phosphate species can stabilize mTOR, 2. inositol phosphates do not detectably change mTORC1 K_M_ for ATP or peptide substrate and 3. mTOR can dynamically sense IP_6_ concentration, at least in the conditions of *in vitro* kinase reactions tested herein.

## DISCUSSION

IP_6_ was proposed to function as a structural co-factor to aid on mTOR folding when other regulatory subunits are absent. Here we show evidence that indicates that mTOR active state is prolonged by IP_4_, IP_5_ and IP_6_ in a dose-dependent and saturable manner, which is more consistent with inositol phosphates having a regulatory role in mTOR stability. Supporting our model, we found that IP_6_/mTOR interaction was dynamic (**Fig 8**). Further, the position of the I-site within the FAT domain is solvent accessible and in crystallographic studies, the I-site is only partially occupied by IP_6_, suggesting IP_6_ can be exchanged with the environment without complete loss of proper folding.

mTOR is a large protein with a C-terminal catalytic domain next to the FRB and FAT (Frap, ATP and TRRAP) domains. The FAT domain of mTOR forms a C-shaped solenoid structure that surrounds the N- and C-lobes of the kinase domain. The N-terminal half of mTOR contains HEAT (huntingtin, EF3A, ATM, TOR) repeats that fold into structures described by Mayer’s group as horn and bridge (Aylett et al., 2016). In the presence of LST8 and Raptor, the active-site cleft of mTORC1 is blocked by LST8, Raptor and the FRB domain of mTOR, which control substrate accessibility and delivery (Aylett et al., 2016; Yang et al., 2017). In addition to substrate availability, which is regulated by Raptor, activation of mTORC1 signaling *in vivo* involves amino acid-dependent translocation of mTORC1 to the lysosome, where growth factor-activated RHEB/GTP is present. This small GTPase binds to portions of the N-terminal HEAT and FAT domains and allosterically activates mTORC1 by causing conformational changes in these domains that are transmitted through the FAT domain and ultimately results in alignment of the 2 lobes and closing of the catalytic cleft (Yang et al., 2017). As a result, K_cat_ of the enzyme increases without changes in K_m_ for substrate. These structural findings emphasize the important role of the FAT domain in the regulation of mTOR catalytic activity. Interestingly, our studies revealed that IP_4_, IP_5_ and IP_6_ promoted a remarkable increase in active truncated mTOR (lacking the N-terminal HEAT domain) and enhancement of total product formation. As for Rheb, IP_6_ did not change the K_M_ of mTORC1 for peptide or ATP, but rather enhanced total peptide phosphorylation only after prolonged incubations. We speculate that IP_6_ binding to the FAT may improve alignment of the N- and C-lobes, as for RHEB.

Although we report a role for inositol phosphates on mTOR kinase regulation *in vitro*, we did not address whether the effects of these small molecules are through binding to the I-site or whether there are additional binding sites for inositol phosphates within the mTOR molecule. We also don’t know whether IP_4_, IP_5_ or IP_6_ can regulate full length mTOR *in vitro* or *in vivo*. The Mayer’s group argued that IP_6_ is dispensable for mTORC2 regulation *in vivo* based on the observation that mTORC2 signal is not affected by knockout or knockdown of the two enzymes that regulate IP_6_, the inositol polyphosphate phosphatase 1 (MINPP1) and the IP5 2-kinase (IPPK) (Scaiola et al., 2020). One potential explanation for this negative outcome is that IP_5_ or IP_4_ can compensate for the loss of IP_6_ to maintain mTOR active, as indicated by the data presented here. In fact, inositol-1,3,4,5,6-P_5_ (IP_5_) concentrations are in the range of IP_6_ in many cells, although its abundance was found to be less consistent than IP_6_ (Qiu et al., 2020). The IP_4_ isomers, inositol-1,3,4,6-P_4_, inositol-1,3,4,5-P_4_ and inositol-3,4,5,6-P_4_ were also detectable through CE-EIMS in human cell lines but at lower levels than IP_5_ and IP_6_. Future experiments will address whether mTORC1 and/or mTORC2 is/are affected in cells lacking IP_4_, IP_5_ and IP_6_ and whether fluctuations in the levels of free inositol phosphates can contribute to physiological regulation of these kinase complexes in specific subcellular compartments.

## MATERIALS AND METHODS

### Compounds

Inositol phosphate and other compounds: Inositol-1-monophosphate (IP_1_), inositol-2,4-bisphosphate (IP_2_), inositol-1,4,5-trisphosphate (IP_3_), inositol-1,3,4,5-tetrakisphosphate (IP_4_ designated isomer A), inositol-1,3,4,6-tetrakisphosphate (IP_4_ designated isomer B), inositol-1,4,5,6-tetrakisphosphate (IP_4_ designated isomer C or simply IP_4_), inositol-1,3,4,5,6-pentakisphosphate (IP_5_), IP_6_ were purchase from Cayman Chemical. Please note that 3 different IP_4_ isofomers were used and distinguished by the letters A, B and C. IP_4_ without any letter indicates the inositol-1,4,5,6-tetrakisphosphate isomer. Inositol, inositol IS_6_ and glucose-6-phosphate were from Sigma.

### mTOR proteins

Recombinant SF21/SF9 expressed and purified FLAG-tagged mTOR (1362-C-term) and mTORC1 [FLAG-mTOR (1362-C-term), HIS-LST8 and HIS-Raptor] proteins were obtained from Milipore/Sigma by Eurofins DiscoverX products (France). We confirmed the presence of HIS-LST8 and HIS-Raptor in the mTORC1 preparations with Raptor less abundant than LST8.

### Kinase assays

Unless indicated, all kinase reactions were in Tris pH 7.5 (50 mM), NaCl (100 mM), MnCl_2_ (10 mM), DTT (1 mM), ATP (10μM) with or without [^32^P]-ATP (approximately 1 μCi/10 μl). Reactions were typically carried out at 30°C for 0.5-1.5 hrs, unless indicated. For autokinase reactions mTOR or mTORC1 were used at 100 ng per reaction (10 ng/μl) and reaction was stopped with 10-15 mM EDTA (unless indicated as in figure 7), before addition of SDS-loading buffer and SDS-polyacrylamide gel electrophoresis (PAGE). Gels were stained with Coomassie stain InstantBlue (Abcam), dried before exposure and analysed using a Typhoon (Cytiva) phosphorimager. For the experiment shown on Supplemental Figure 1, Coomassie stained bands were visualized and analyzed using Odyssey (LiCor) infra-red scanner and Image Studio software. Peptide kinase assays were designed based on the work by Sabatini’s group (Kang et al., 2013)with modifications and using the following peptides as substrate (HPLC purified, Biomatik): GYDYSTTPGGTGRRRRR (derived from 4EBP S65 phospho-site) and GYFLGFTYVAPGRRRRR (derived from p70S6K T389 phospho-site). Unless otherwise indicated, 4EBP-derived peptide was used at 40-100 μM and mTOR was used at 5-10 ng/μl or 33-66 nM. Reactions were stopped with EDTA (15 mM) and spotted in triplicates into W3 cation exchange filter paper (from Jon Oakhill, St. Vincent’s Institute Medical Research, Australia). Filters were washed in 0.42% H_3_PO_4_, for three 30-minute cycles and left in 0.42% H_3_PO_4_ overnight before air drying. A mixture of buffer and [^32^P]-ATP was spotted as negative control for background detection. The radioactivity present in each spot was quantified using phosphorimager (Typhoon, Cytiva) and Image Studio software or by mixing each spot with scintillation fluid and measuring cpm with scintillation counter (Packard Tricarb, PerkinElmer and Quantasmart software). For the phosphatase inhibitor experiments (Figure 3), NaFl (0.5 mg/ml), ß-glycerophosphate (0.5 mg/ml) and nor-cantharidin (5 μM) were used together with protease inhibitor cocktail (Pierce).

### Western blots and solubility assay

Solubility assays were set up exactly as for the kinase assays using peptide as substrate with the exception that only cold ATP was used. mTOR concentration was 5 ng/μl and was diluted in buffer containing protease inhibitors. After incubation at 30°C, 3 separate samples of the supernatant (2.5 μl each) were collected and transferred to a new tube containing EDTA (15 mM) and SDS-gel loading buffer. The mTOR leftover in the tubes were extracted with EDTA/SDS loading buffer and considered the insoluble fraction. All samples were boiled and separated on SDS-PAGE using a pre-cast gradient gel (4-15%, BioRad). Proteins were transferred to nitrocellulose membrane, which was then blocked with 5% milk and blotted using anti-mTOR antibody (Cell Signaling Technologies) as primary antibody. Anti-rabbit conjugated with infra-red dye 680 was used as secondary antibody and membranes were scanned using the Odyssey infra-red scanner (LiCor) and analyzed with Image Studio software. For the experiment shown in Figure 4 A and B, triplicate reactions were set up separately and 2.5 μl samples were collected for each time point which is equivalent to ¼ of the total volume. For the experiment shown in Figure 4 C and D, kinase and solubility reactions were run in parallel.

### Gel shift assays

mTOR electrophoretic mobility shift was assayed as for autokinase assay with [^32^P]-ATP or cold ATP. Reactions were set up exactly as for kinase reactions, unless indicated otherwise. However, for the super-shift assay, SDS-loading buffer was used to stop the reactions without any EDTA being present. When [^32^P]-ATP was used, mTOR was detected using phosphorimager (Typhhon, Cytiva) and when cold ATP (or no ATP) was used, mTOR was detected using western-blot as described above. Typically, less mTOR per lane was used when detection was through western-blot (5-10 ng/lane).

## ACKNOWLEDGMENTS

This work was supported by the National Institute on Aging, R21 NIA AG071975 to L.E.R and R.D.B., National Institutes of General Medical Sciences R01 GM132592 to R.D.B and R01 GM124404 to J.Y. and L.E.R.

## SUPPLEMENTAL FIGURES

**Supplemental Figure 1:**
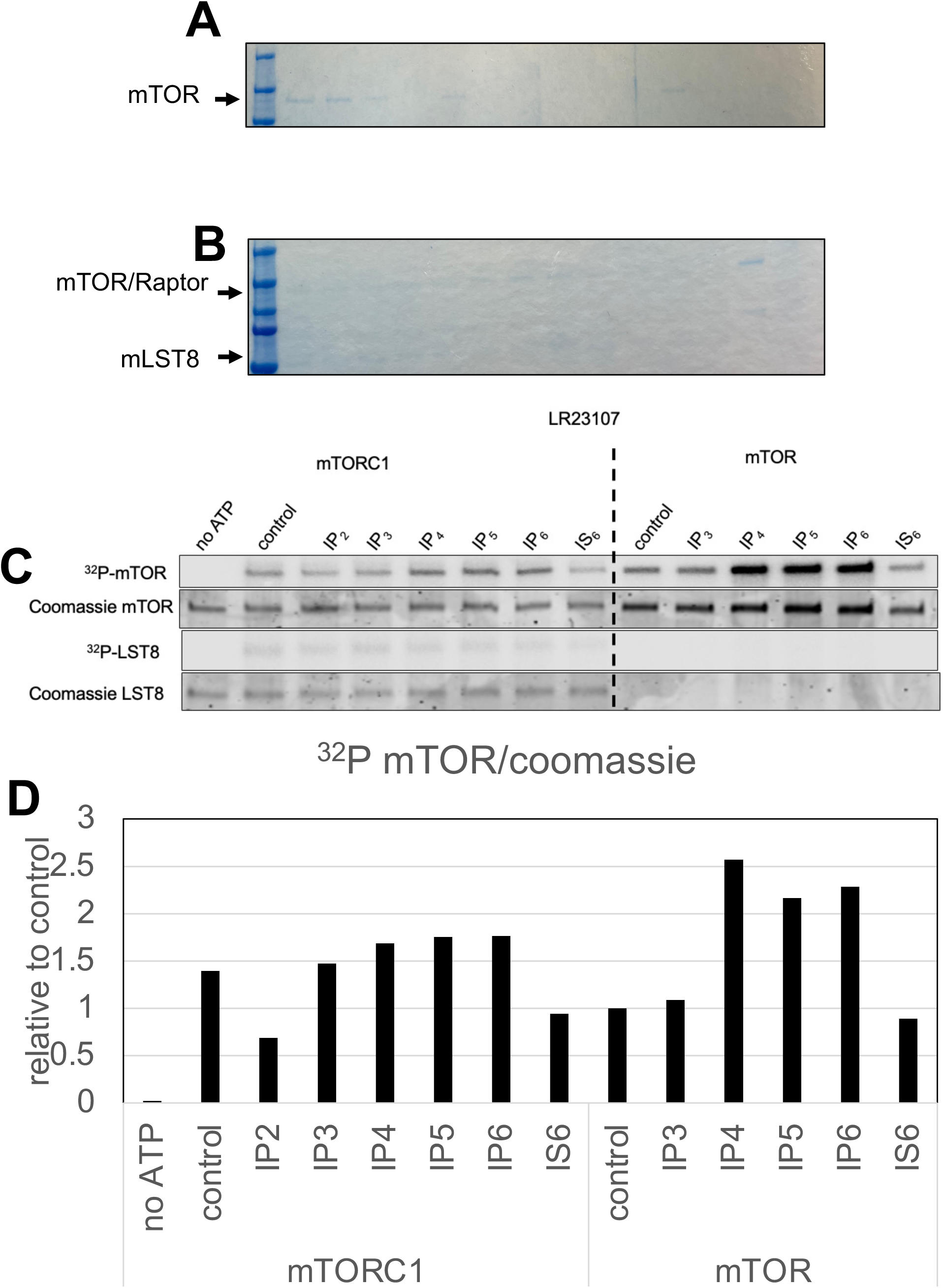
Coomassie stain images of the gels shown on Figure 1A (A) or 1B (B). Panel C shows the phosphorimager image and infra-red image of the equivalent Coomassie stained gel (C) with quantification and normalization of the counts using Coomassie stained mTOR (D). Also shown is [^32^P]-LST8 and coomassie stained LST8. In this experiment a total of 200 ng mTOR was used per lane for better quantification of the Coomassie bands.

**Supplemental Figure 2:**
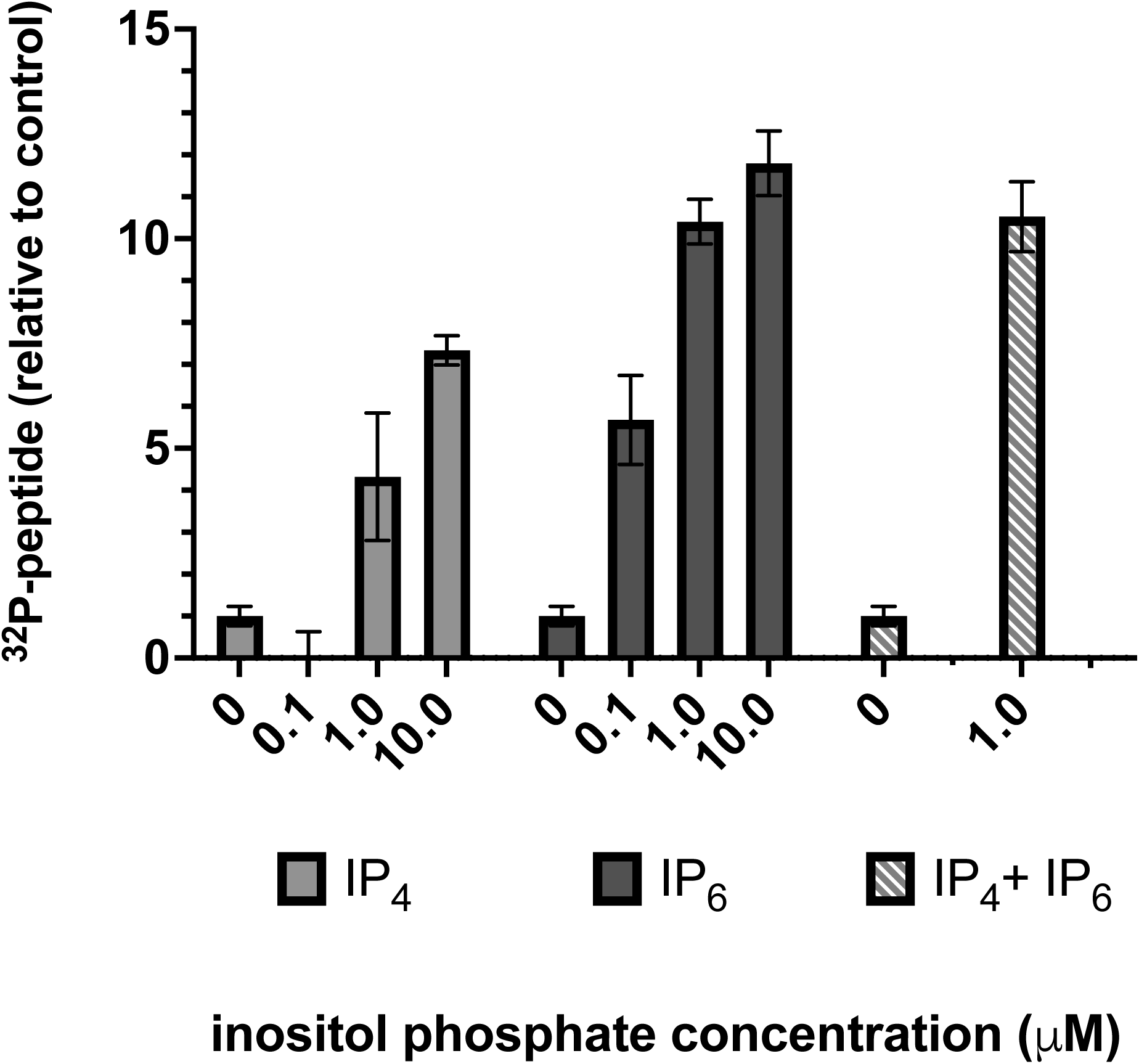
IP_4_ and IP_6_ enhance mTOR phosphorylation of peptide substrate with different affinities and in a non-additive manner. mTOR was incubated with [^32^P] γ-ATP in kinase reaction without (control) or with 0.1, 1 or 10 μM of IP_4_ or IP_6_, as indicated or with 1.0 μM of plus 1.0 μM IP_6_ (hatched bars). Reactions were stopped with EDTA before spotting on P81 paper. Results shown is the mean and standard deviations of quantified triplicate spots.

**Supplemental Figure 3:**
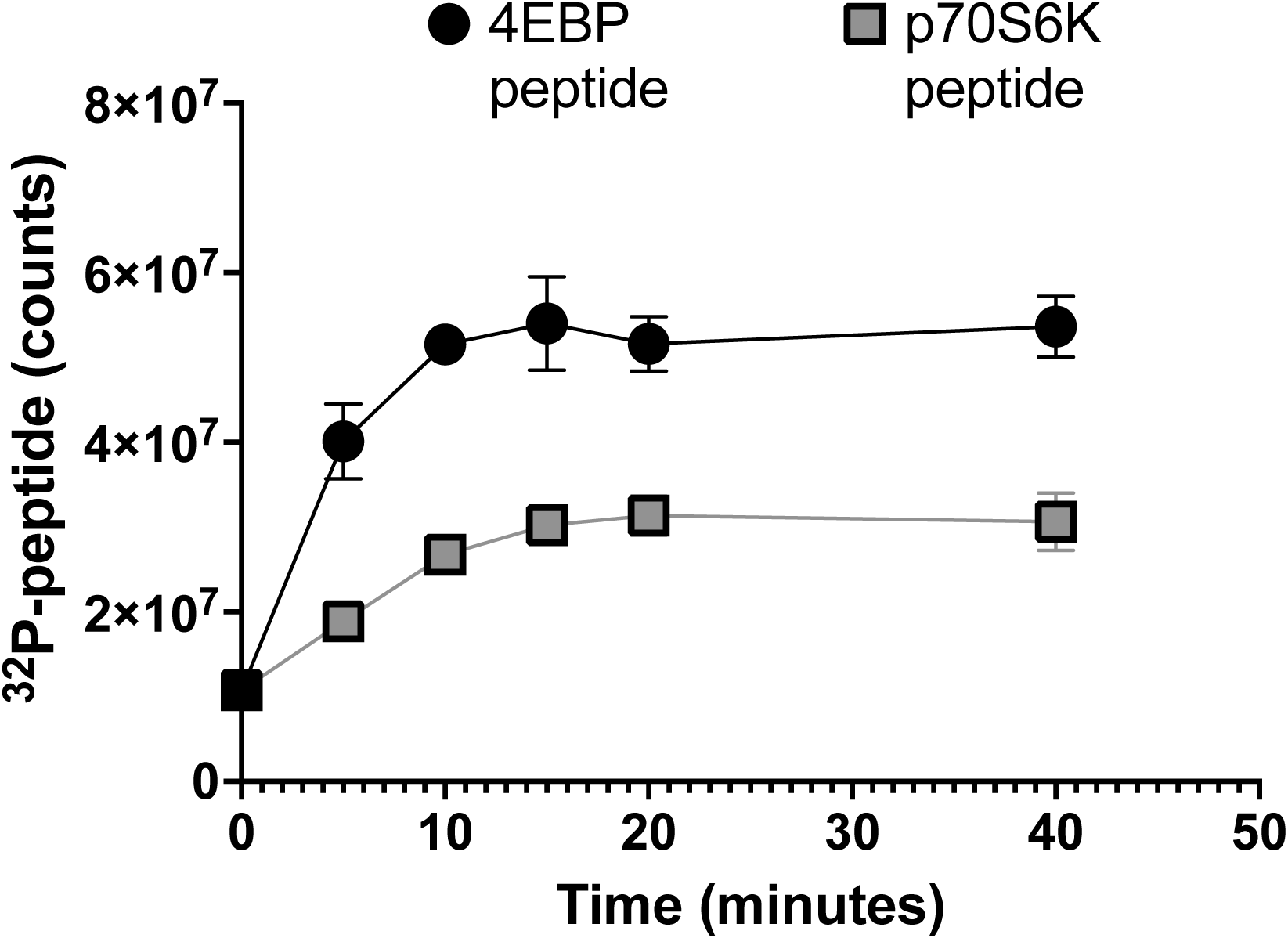
mTOR kinase towards 4EBP or p70S6K peptide substrate is short lived. mTOR kinase activity was assay using 4EBP or p70S6K-derived peptide substrate with [^32^P] γ-ATP in kinase reaction without any inositol phosphate. Samples were collected at the time indicated for spotting on P81 paper and analysis of the counts present in the peptides.

**Supplemental Figure 4:**
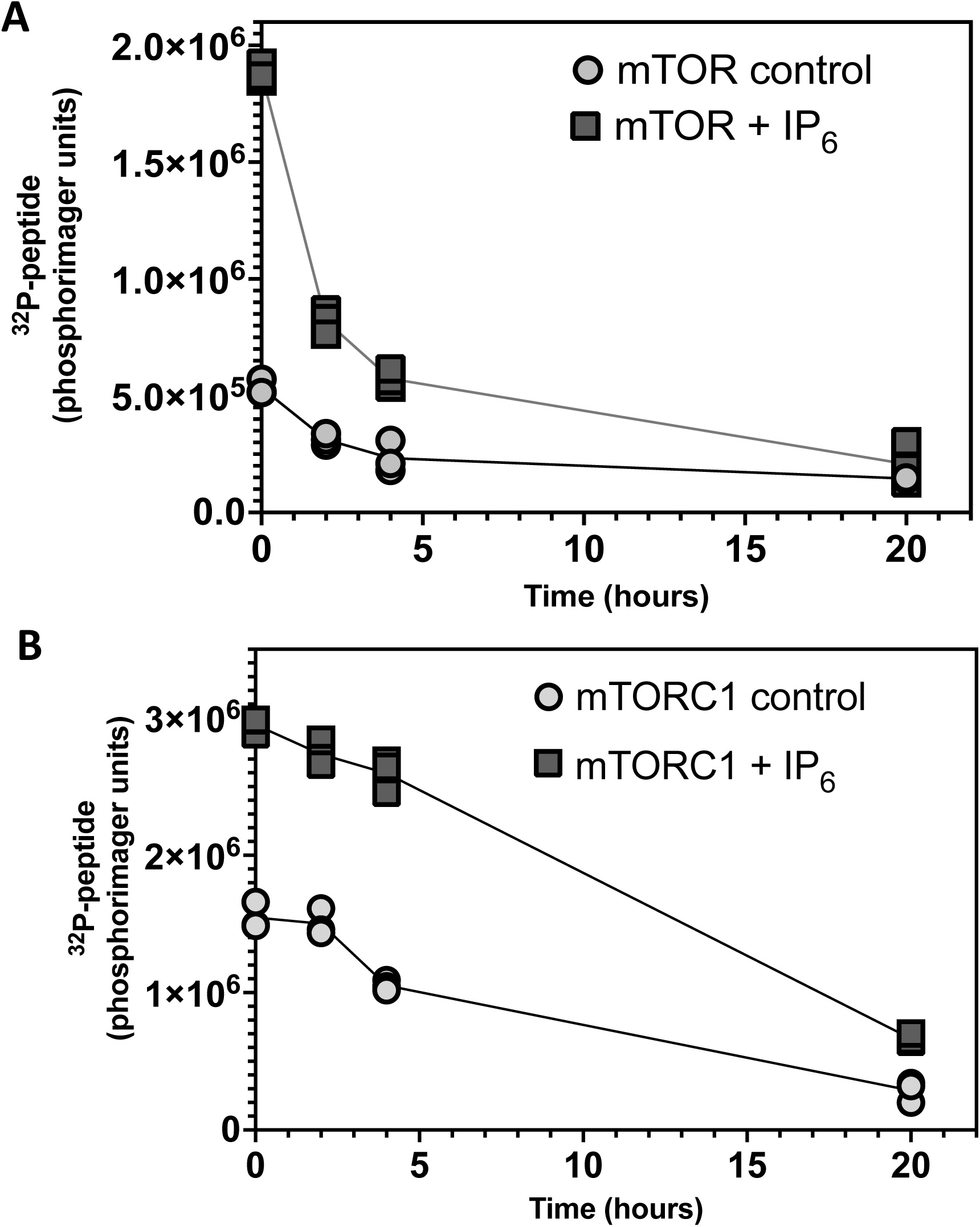
mTOR (A) and mTOR/LST8/Raptor (B) stability over time. mTOR (A) or mTOR/LST8/Raptor (B) were incubated without or with 10 μM IP_6_ at room temperature in kinase buffer containing all components except ATP. At the time indicated, samples were collected, [^32^P]-ATP/ATP was added and assayed for 30 minutes at 30°C. Data shown are the mean and standard deviations of triplicate samples spotted on P81 paper and counted in phosphorimager.

**Supplemental Figure 5:**
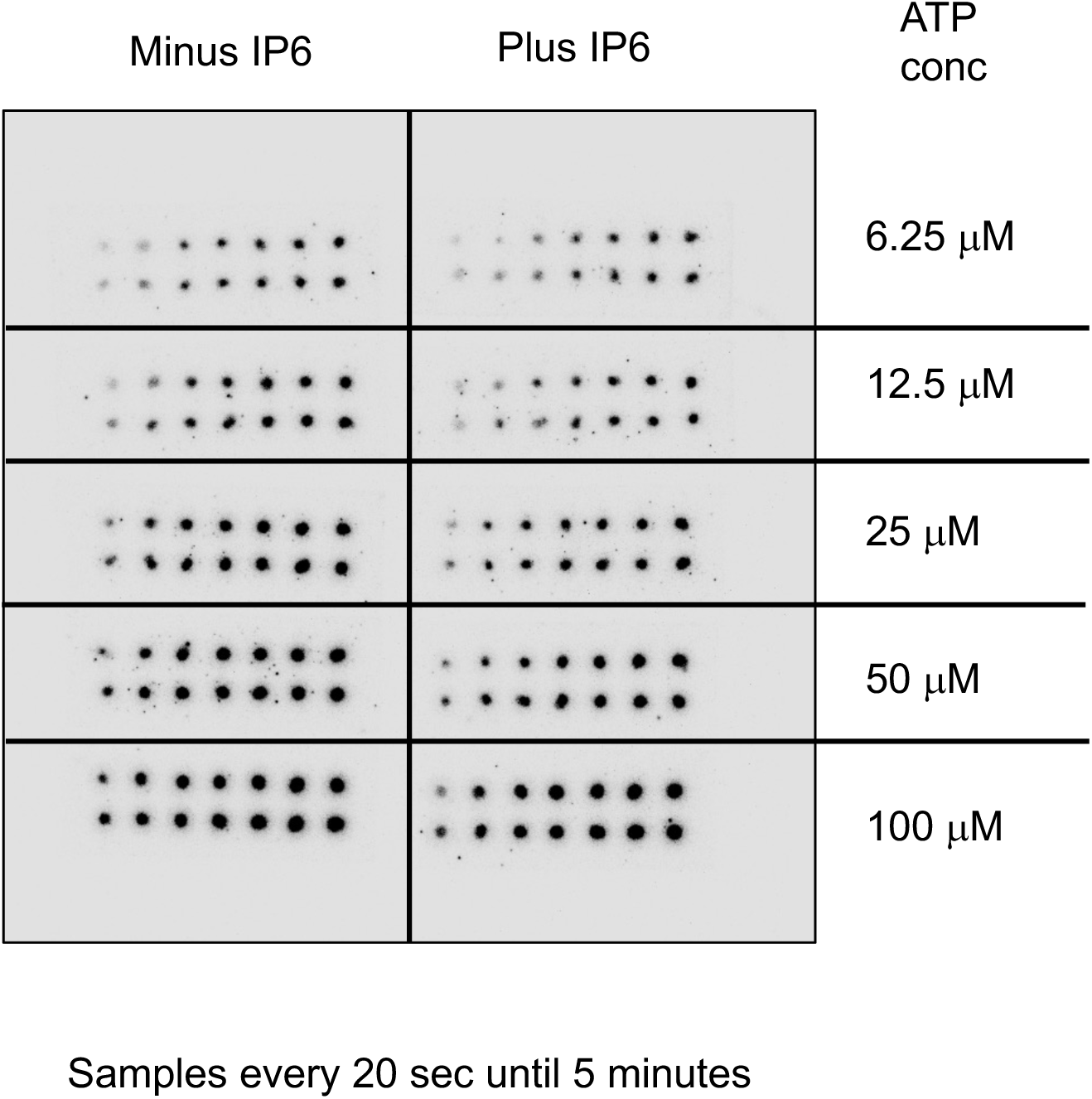

